# Complement 3 signaling is necessary for the developmental refinement of olfactory bulb circuitry

**DOI:** 10.1101/449454

**Authors:** Katherine S. Lehmann, Alynda Wood, Diana Cummings, Li Bai, Beth Stevens, Leonardo Belluscio

## Abstract

The olfactory system depends upon organizational maps that are developmentally refined and maintained, however the cellular and molecular mechanisms that underlie these processes are unknown. Studies have shown that microglia and complement molecules are important for the developmental refinement of circuitry within the visual system, thus we asked whether they played a similar role in the olfactory system through the formation of the olfactory bulb (OB) maps, the glomerular and intrabulbar maps. Our findings revealed that microglia in mature animals engulf olfactory sensory neuron (OSN) axons and the synaptic terminals of tufted cells in the glomerular and intrabulbar maps respectively, suggesting microglia could anatomically shape the mature OB circuitry. To determine the mechanisms underlying this axonal pruning activity we used complement 3 (C3) and complement receptor 3 (CR3) knockout mice to investigate if C3 signaling was necessary for precise OB map development. Our results demonstrate that glomerular and intrabulbar map disorganization as typically present in early postnatal mice persists into adulthood when C3 signaling is disrupted. These data clearly establish the C3/CR3 pathway as necessary for the proper developmental refinement of both olfactory maps. We further present the olfactory system as a unique platform to study the role of glia in the development and adult refinement of regenerating circuits.

## Introduction

The olfactory system depends upon organizational maps that are established perinatally and refine to adult precision during maturation with varying dependence on sensory activity (Zou *et al.*, 2004; Marks *et al.*, 2006). The mechanisms that underlie the refinement process remain unknown.

The glomerular map is composed of olfactory sensory neurons (OSNs), a regenerating neuronal population in the olfactory epithelium (OE) that send axons to the surface of the olfactory bulb (OB), where they converge to form odorant receptor (OR)-specific glomeruli (Graziadei *et al.*, 1978; Costanzo, 1991; Belluscio *et al.*, 2002; Kerr & Belluscio, 2006). Developmental studies have shown that during embryonic olfactory bulb formation OSN axons are often mistargeted either to the wrong glomeruli or to the incorrect layer of the olfactory bulb (Royal & Key, 1999; Cho *et al.*, 2007). Evidence of mistargeted axons persists into the early postnatal period forming smaller atypical glomeruli or “minor” glomeruli which largely disappear by 4 weeks of age (Zou *et al.*, 2004), Thus by adulthood, incorrectly targeted OSN axons are rarely observed.

The intrabulbar map undergoes a similar refinement process as its broad projections gradually refine to link isofunctional glomeruli during its postnatal maturation. In the mouse OB each glomerulus has a “twin” on the opposite side of the OB and the intrabulbar circuitry allows them to interact (Lodovichi *et al.*, 2003). This map is mediated by tufted cells, which receive glomerular input from one side of the OB and extend their axons to the internal plexiform layer (IPL) on the opposite side of the OB forming synaptic connections with interneurons just below the twin glomerulus on the opposite side (Cummings & Belluscio, 2010). Tufted cell intrabulbar projections undergo both developmental and activity dependent refinement; their axons broaden in times of sensory deprivation and are refined when activity is restored (Marks *et al.*, 2006; Cummings & Belluscio, 2010). The mechanism of both the developmental and sensory dependent refinement is still unknown and although the neurons that mediate these maps have been characterized the involvement of other cell types in this process is also unclear.

Recent studies suggest that microglia may be responsible for shaping neuronal circuitry during development (Nimmerjahn *et al.*, 2005; Stevens *et al.*, 2007; Paolicelli *et al.*, 2011; Schafer *et al.*, 2012; Sipe *et al.*, 2016) and disrupting circuitry linked to some neurological disease (Frick & Pittenger, 2016; Hong *et al.*, 2016; Vasek *et al.*, 2016; Williams *et al.*, 2016). This raises the possibility that microglia play a significant role in circuitry establishment and refinement throughout an organism’s lifetime. These studies identified the complement cascade and in particular complement 3 (C3) signaling as a key mechanism for synaptic pruning in the visual system (Stevens *et al.*, 2007; Schafer *et al.*, 2012) and hippocampus (Shi *et al.*, 2015), indicating that C3 signaling may be broadly necessary for developmental circuitry refinement.

Here we investigate the role of microglia in olfactory bulb circuitry development and refinement. We test whether complement signaling is responsible for proper olfactory development by examining two different circuits in the olfactory bulb. We used transgenic mice to visualize microglia, OSN axons and tufted cells and to knockout C3 signaling. Additionally, we applied a tetramethylrhodamine (TMR) tracer injection assay that allows for direct visualization and measure of intrabulbar projections (Marks *et al.*, 2006; Lodovichi & Belluscio, 2012). The data show microglia engulfment of axonal and dendritic processes in mature animals, a novel finding, and that C3 signaling is necessary for the developmental refinement of both OSN axons and tufted cell projections, providing a new mechanism of developmental circuit refinement in the OB.

## Materials and Methods

### Experimental animals

All animal procedures complied with National Institutes of Health guidelines and were approved by the National Institute of Neurological Disorders and Stroke Animal Care and Use Committee. Mice were kept on a 12 h light/dark cycle with food and water *ad libitum*.

### Transgenic Lines

We used C3(-/-) animals (https://www.jax.org/strain/003641) where the C3 gene is disrupted by insertion of a PGK/Neo cassette and CR3(-/-) where the CR3 gene is disrupted by vector insertion in the translation initiation codon (https://www.jax.org/strain/003991). AP-2µ-Cre mice were obtained from Trevor Williams at the University of Colorado (https://www.ncbi.nlm.nih.gov/pmc/articles/PMC2745980/). P2-IRES-tauGFP (https://www.jax.org/strain/006669) mice, Rosa-tdTomato (https://www.jax.org/strain/007905) Rosa-tdTomato-Synaptophysin (https://www.jax.org/strain/012570) and Cx3Cr1 (https://www.jax.org/strain/005582) were obtained from The Jackson Laboratory

### Transcardial perfusion and immunohistochemistry

Mice were deeply anesthetized with ketamine (300 mg/kg) and xylazine (6 mg/kg) and perfused transcardially with 0.1 M PBS, then 4% PFA. OBs were post-fixed overnight (4% PFA), cryoprotected (30% sucrose), embedded (10% gelatin), fixed/cryopreserved (15% sucrose/2% PFA) overnight and sectioned coronally at 45 μm. The cutting angle was carefully adjusted to ensure that sections contained equivalent coronal planes through the left and right OBs. For immunohistochemistry, sections were blocked for 1 h (5% horse serum/0.5% Triton X-100 in TBS), followed by 48 h with primary antibody on a shaker (4°C) and 2 h with secondary antibody (room temperature). Sections were mounted with Vectashield H-1200 with DAPI (Vector Laboratories). Antibodies were IBA1 (Wako, #019-19741) and CD68 (Bio-Rad, #MCA1957).

### Immunofluorescence image acquisition and analysis

Images were acquired using a Leica TCS SP5 microscope. GFP fluorescence was collected at 500–550 nm and tdTomato fluorescence at 600–650 nm. Widefield images were acquired using an HCX PLAN FLUOTAR 10x/0.30 NA air objective. Confocal images were acquired using an Achroplan 20x/0.45 objective and an HCX PL APO 40x/1.25 NA oil-immersion objective. CD68 puncta and Iba1 area quantification was automated using Fiji. P2-GFP glomeruli were counted and classified manually.

### Tracer injections

Mice were anesthetized with Ketamine (200 mg/kg, ip) and Xylazine (10 mg/kg, ip) and maintained with isofluorane (1%-3% in 100% O_2_). The scalp was resected and the bone over the olfactory bulbs was removed. Tracer injections of 10% dextran tetramethylrhodamine (TMR, 3000 mw) were targeted to the glomerular layer and iontophoresed through a micropipette (10 µm tip diameter). At 12-24 hr post-injection animals were transcardially perfused with 1X PBS, followed by 4% PFA. Brains were removed and post-fixed in 4% PFA, cryoprotected in 30% sucrose, microtome sectioned (60 µm horizontal sections) and mounted onto slides (Vectashield).

### Tracer image acquisition and analysis

Images of tracer injection and projection sites were collected using a Zeiss LSM 510 laser scanning confocal microscope at both 10× and 20× magnifications (excitation 554, emission 580). The diameter of the projection site was defined as the longest distance between branch points of labeled axons using a line drawn parallel to the mitral cell layer. Ratios were calculated between the diameters of the injection site and corresponding projection site, averaged, and reported ±SEM. Comparisons of averaged projection: injection site ratios were analyzed with a one-way ANOVA followed by *post hoc* comparisons using the Holm–Sidak method.

### Statistics

Statistical analysis was performed in Prism (Graph Pad 7). Data that were normally distributed with equal variance were analyzed using student *t*-test or one-way ANOVA. Post-hoc analysis is indicated when used. All values are reported as the mean ± SEM. The *p* values (\**p* < 0.05 and \*\**p* < 0.001) are also indicated in the corresponding figure legend. n=animal number.

## Results

### Microglia engulf OSN axons

Glomerular map organization is established during development and refined through activity dependent pruning of unorganized or mistargeted axons (Zou *et al.*, 2004; Kerr & Belluscio, 2006). Once the map is established it is maintained despite the constant regeneration and innervation of new axons (Costanzo, 1991). To determine if microglia participate in glomerular map refinement we first looked for evidence of microglia engulfment of OSN axons in the glomerular layer. We used 8-week old adult homozygous P2-GFP animals to specifically label olfactory receptor 17 (Olfr17) glomeruli (Royal & Key, 1999; Feinstein & Mombaerts, 2004). At this age, animals will typically have two isofunctional, fully formed glomeruli representing each OR, which we termed “major glomeruli” (Figure 1A) (Royal & Key, 1999). Additionally, there are often bundles of mistargeted axons that converge but do not form a full glomerular structure, “minor glomeruli” (Royal & Key, 1999; Zou et al., 2004) (Figure 1B). We stained sections containing both types of structures with Iba1, a microglia maker, to visualize the association of microglia with glomerular axons. This revealed axonal engulfment around both major glomeruli, and minor glomeruli and we often observed GFP positive particles inside the microglia around the glomeruli (Figure 1C). A 3D orthogonal view provided further evidence of engulfment (Figure 1D) and confirmed that microglia can engulf OSN axons.

**Figure 1.**
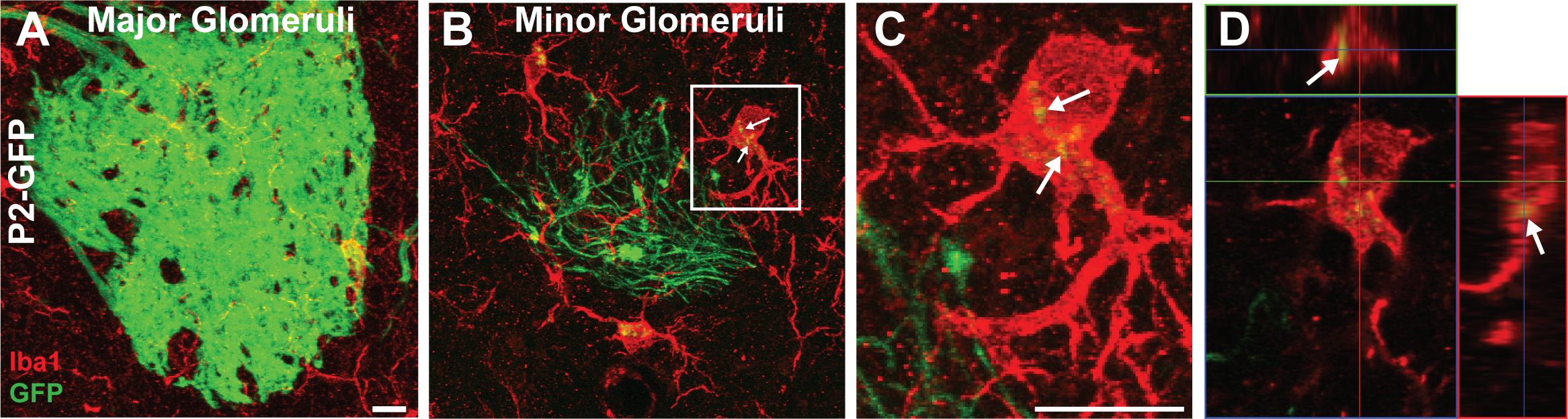
Microglia engulf olfactory sensory neurons in adult mice. **(A)** Immunofluorescence staining on a major, stereotypical P2-GFP OSN glomeruli against the Iba1 (red) microglia marker in the olfactory bulb of an 8 week old mouse **(B)** Representative minor glomeruli made of mistargeted P2-GFP OSNs stained with Iba1 (red). **(C)** High magnification image of a microglia with engulfed P2-GFP axons internalized, indicated by arrows. **(D)** Same microglia in an orthogonal 3-D view of microglial engulfment. Scale bar =10μm.

### Microglia engulf tufted cell synaptic terminals

The intrabulbar map, composed of tufted cells, is also subject to activity dependent refinement, and like the glomerular map is developmentally refined (Marks *et al.*, 2006). However, unlike the glomerular map, tufted cells do not degenerate and regenerate (Cummings & Belluscio, 2010) and therefore allow us to assess how microglia interact with a stable population of neurons in the adult brain.

To study tufted cells, we used ten week old animals from a transgenic AP-2ε-Cre line (Feng *et al.*, 2009), a tufted cell specific driver, crossed with Rosa-tdTomato-loxP animals (Figure 2A). We specifically focused on tufted cell dendrites in the glomerular layer (Figure 2B) and tufted cell axons in the internal plexiform layer (IPL) (Figure 2C) because these processes make dendrodendritic and axonal dendritic connections respectively. To address if microglial engulfment of these processes could shape intrabulbar map circuitry through synaptic pruning we crossed the AP-2ε-Cre animals with a recombinase dependent synaptic reporter line, Rosa-tdTomato-Synaptophysin, and then crossed those with CX3CR1^GFP/GFP^ animals to create a line with fluorescently labeled tufted cell synapses and fluorescently labeled microglia (Figure 2D). We found that microglia in both layers engulfed synaptic terminals (Figure 2D), evidenced by synaptophysin positive debris in the microglia. We focused on the IPL layer (Figure 2E), the location of the intrabulbar map and found microglia internalized Synaptophysin (Figure 2F, G), indicating that microglia engulf synapses in the olfactory bulb.

**Figure 2.**
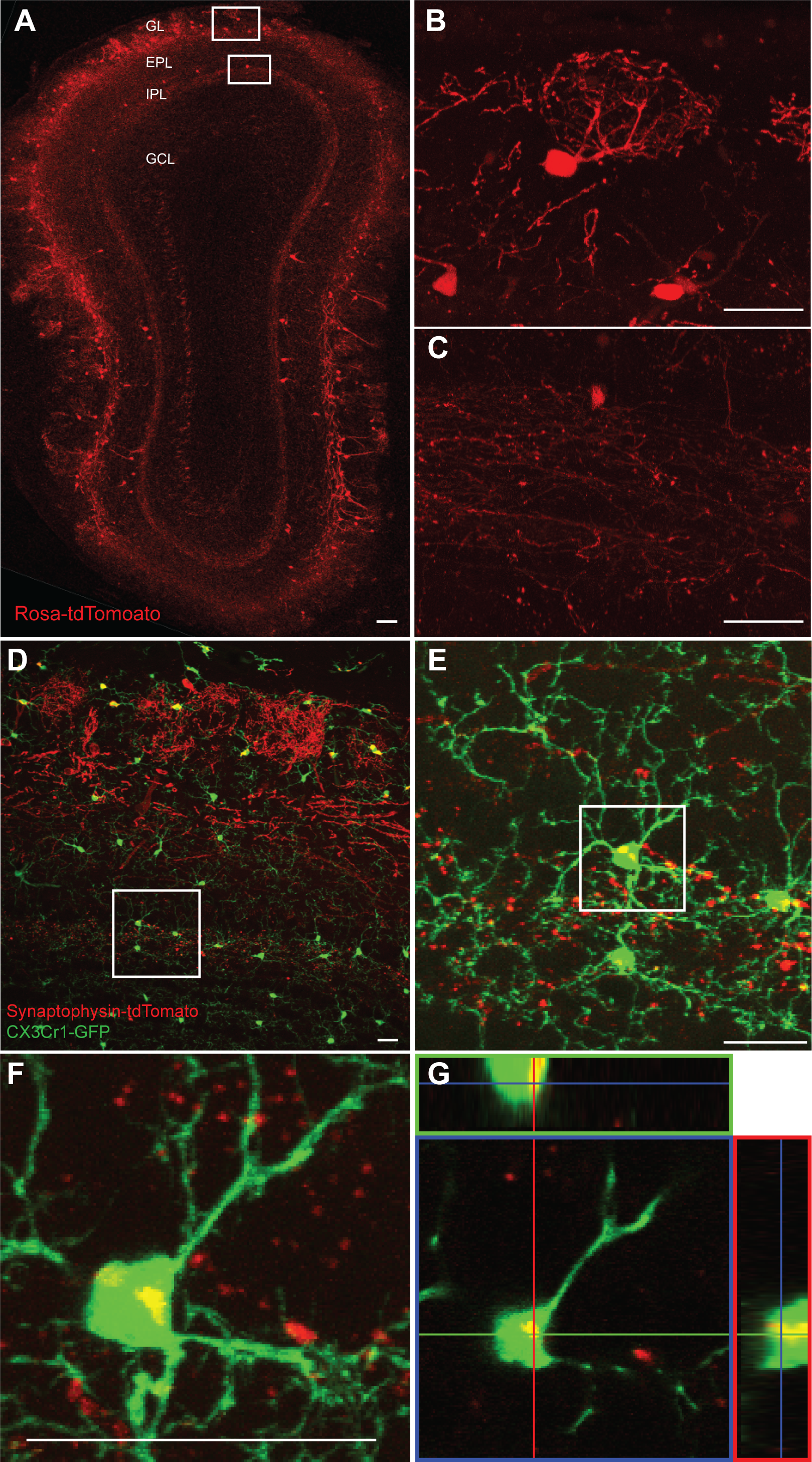
Microglia engulf tufted cell axons and dendrites. **(A)** Coronal section of the olfactory bulb. Tufted cells (red) labeled with AP2e-Cre; Rosa-tdTomato-loxP reporter line showing layers of the bulb: Glomerular Layer (GL), External plexiform layer (EPL), Internal plexiform layer (IPL) and Granule cell layer (GCL). **(B)** High magnification tufted cell soma and dendrites in the glomerular layer and **(C)** tufted cell axons in the IPL. **(D)** Coronal section of olfactory bulb with tufted cell synaptic terminals (red) labeled transgenically with the Ap2e-Synaptophysin-TdTomato, and microglia (green) labeled via the CX3CR1-GFP transgenic line. **(E)** Microglia in the IPL with engulfed tufted cell synaptic terminals internalized **(F, G)** High magnification images of microglia engulfment of tufted cell synaptic terminals in the IPL (arrows in C, D). Shown as **(F)** Z-projection and **(G)** orthogonal 3-D view Scale bar A=100μm Scale bar B-F =25μm.

### Microglia lysosomal activity in the IPL of the olfactory bulb is dependent on C3

Previous studies propose the complement 3 pathway, specifically the complement 3 receptor (CR3) and its C3 ligand as important for microglia activity, and for microglial mediated refinement (Sierra *et al.*, 2010; Tremblay *et al.*, 2010; Schafer *et al.*, 2012; Shi *et al.*, 2015). To directly test whether complement 3 is important for microglia engulfment activity in the olfactory bulb we employed C3(-/-) and CR3(-/-) animals that remove the C3 ligand and receptor respectively and then stained with CD68, a microglia lysosomal marker. For this analysis we focused on the IPL where we evaluated synaptic engulfment of tufted cell axons. Tissue from the IPL layer in C3 knockout animals showed no reduction in the area of tissue covered by microglia, measured by the percent of area Iba1 positive cells (7.269 ± 1.204, n=14, 6.557 ± 0.7802, n=3: p=.6096) (Figure 3A, B). However, these images showed a drastic decrease in microglia activity, measured as the number of CD68 positive puncta (48.86 ± 4.63, n=14, 26.81 ± 3.988, n=3 p=.0011) (Figure 3C, D). Thus, eliminating C3 signaling had little effect on microglia viability but significantly reduced their activity suggesting that there may also be circuitry refinement deficits in the olfactory bulb with the loss of C3 signaling.

**Figure 3.**
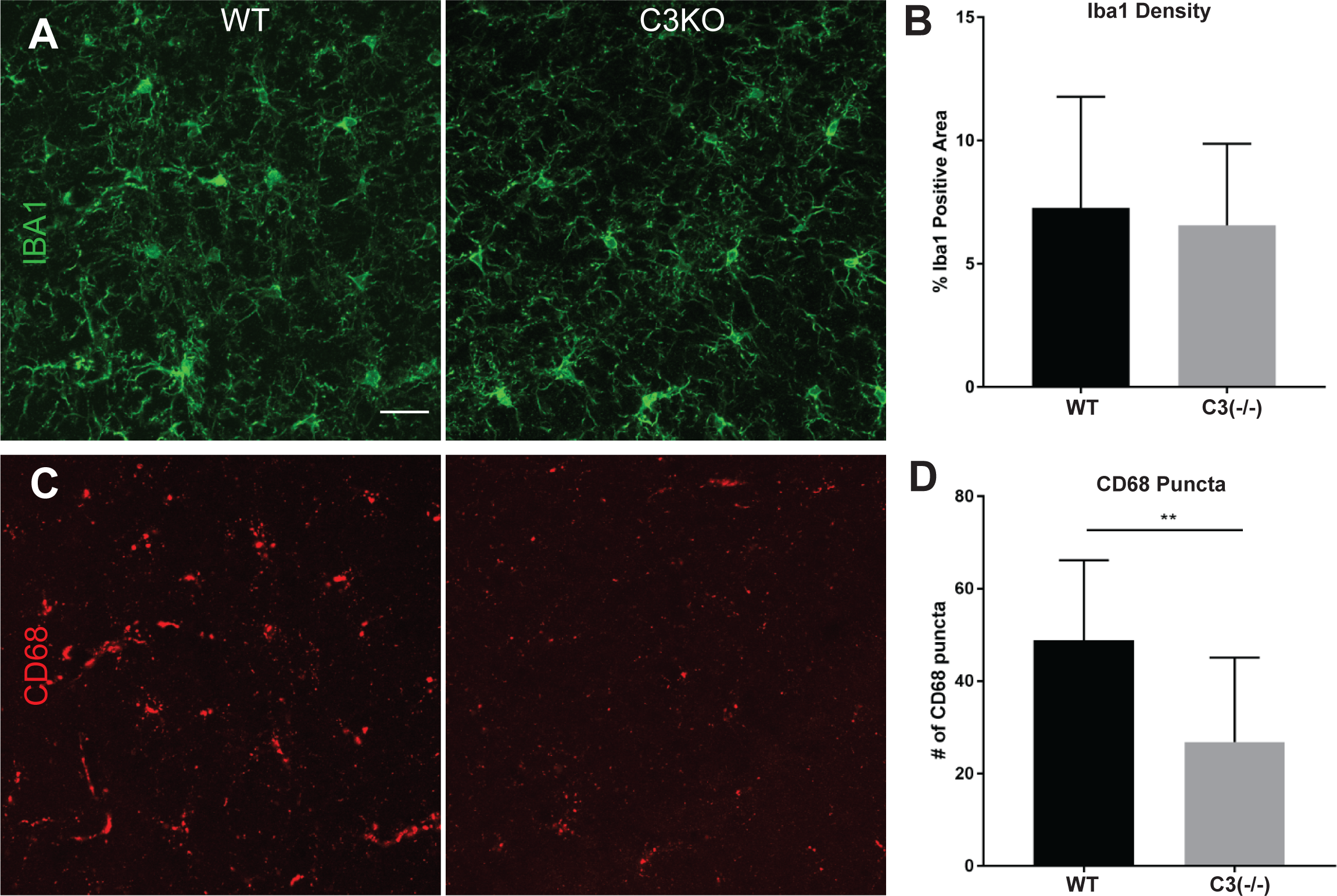
C3 signaling is necessary for microglia lysosomal activity. **(A)** Representative images from the IPL layer stained with Iba1 (green) to mark microglia. **(B)** Percent of area that is Iba1 positive, marking microglia, does not change in C3 knock out animals (7.269 ± 1.204, N=3, 6.557 ± 0.7802, N=3: p=.6096). **(C)** Representative images from the IPL layer showing a decrease in microglia lysosomal marker, CD68 (red) in C3 knockout animals. **(D)** Number of Cd68 puncta in WT and C3(-/-) animals shows significant decrease of Cd68 puncta in C3(-/-) (48.86 ± 4.63, n=3, 26.81 ± 3.988, n=3 p=.0011). Scale bar = 25μm.

### C3 is necessary for tufted cell projection refinement

Given that we saw microglia engulfment of tufted cells and a decrease in IPL microglia activity in C3 mutant mice, we wanted to investigate whether the C3 signaling mechanism was necessary for refining intrabulbar circuitry. To address this question, we used TMR tracer injections (Vasek *et al.*, 2016; Marks *et al.*, 2006) which allowed us to analyze the tufted cell axonal arbor. As an indicator of map refinement, we measured the width of the labeled axonal projection in the IPL and compared it to the width of the injection site in the glomerular layer. The ratio of injection to projection values provides a specificity measure for the intrabulbar projections (Figure 4). In wildtype mice (n=5) we observed an average of 1:1.41 ± .03 injection: projection ratio, similar to previous studies (Figure 4A) (Marks *et al.*, 2006). Interestingly, in C3(-/-) (n=5) (Figure 4B) and CR3(-/-) (n=4) mice we found much wider arbors (Figure 4C), nearly doubling the ratio observed in control animals (n=5, 1:3.20 ± .22; n=4,1:3.46 ±.04). Data analysis using a one-way ANOVA (Figure 4D) revealed significant differences across groups (F2,10 =250.39; P<0.001) and *post hoc* tests showed that the average projection site widths of C3(-/-) and CR3(-/-) mice were significantly broadened compared to controls (p< 0.0001, t=9.502: p<0.0001 t=10.25) Thus, we conclude that C3 signaling is necessary for refinement of intrabulbar tufted cell projections.

**Figure 4.**
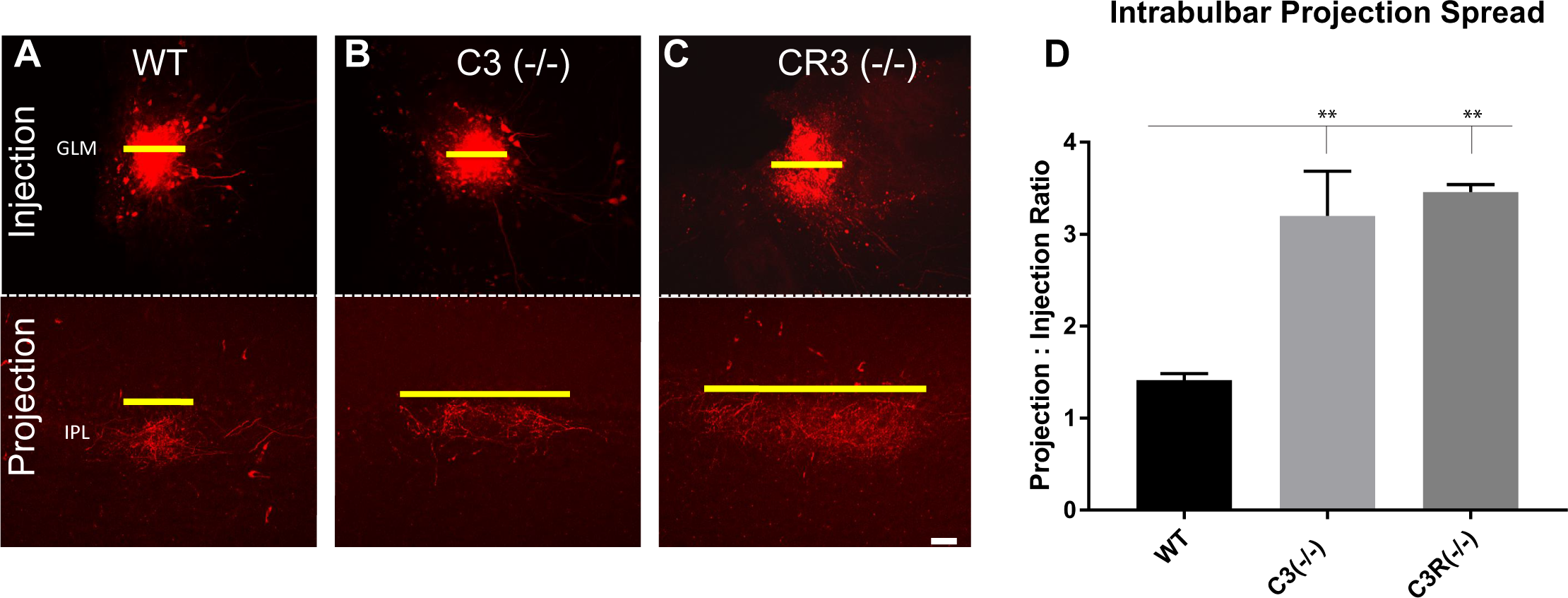
C3 signaling is necessary for pruning intrabulbar projections. TMR tracer injection at the injection site and projection site to visualize intrabulbar projections. **(A)** Wildtype animals (n=5) had an projection:injection diameter ratio 1:1.4. **(B)** C3 (-/-) mice (n=4) TMR tracer injections had an projection:injection diameter ratio of 1:3.2 and **(C)** CR3(-/-) mice (n=4) had a ratio of 1:3.5. **(D)** Data analysis using a one-way ANOVA revealed significant differences across groups and *post hoc* tests showed that the average projection site lengths of C3 and CR3KO mice were significantly broadened compared to controls (p<.0001). Scale bar = 50μm.

### C3 is mechanism of glomerular pruning

We then investigated whether this C3/CR3 dependent refinement was also necessary in the glomerular map. To accomplish this, we made P2-GFP; C3(-/-) and P2-GFP; CR3(-/-) compound mutant animals. We examined 8-10 week old animals and analyzed the large stereotyped major glomeruli as well as the minor glomeruli to determine if C3 signaling played a role in the formation and refinement of these structures. (Figure 5A, B, C). In all cases the number of major glomeruli remained constant (Figure 5D) (F (2, 31) = 0.6444, p=.5319). However, the number of minor glomeruli significantly increased, nearly doubling, across the C3(-/-) (n= 10) and CR3 (-/-) groups (n=6) (F (2, 31) = 36.52, p<.0001) (Figure 5D). Post hoc tests using the Holm—Sidak method to compare wildtype controls to the C3(-/-) and CR3 (-/-) groups found significant increases for both groups (p<.0001 t=8.02: p<.0001, t=5.214). The increase in minor glomeruli when either C3 or the CR3 receptor is knocked out indicates that microglial C3 signaling is necessary for the refinement of mistargeted OSN axons, revealing a new mechanism of glomeruli map refinement.

**Figure 5.**
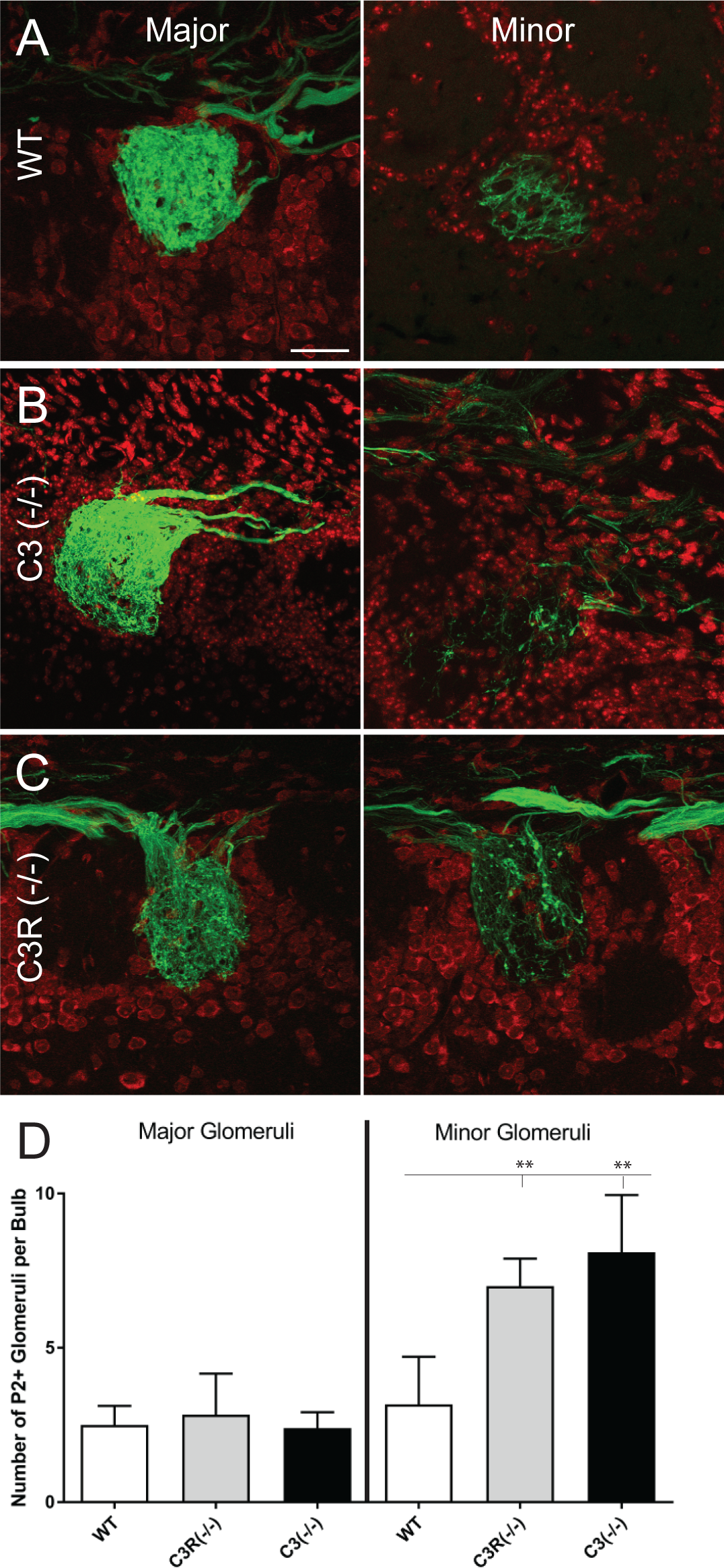
C3 signaling is necessary for pruning axons in minor glomeruli. **(A, B, C)** Representative pictures of P2-GFP major and minor glomeruli in adult (8 week old) animals, counterstained with Nissl (red) in **(A)** Wildtype(n=17), **(B)** C3 (-/-) (n=10) **(C)** CR3 (-/-) (n=6) mice. **(D)** Number of P2-GFP major and minor glomeruli in WT C3(-/-) and CR3(-/-) animals. The number of major glomeruli does not change (F (2, 31) = 0.6444, p=.5319). Post-hoc analysis found a significant increase in the number of minor glomeruli in the C3(-/-) and CR3(-/-) conditions (p<.0001 t=8.02: p<.0001, t=5.214). Scale bar = 25 μm.

## Discussion

We demonstrate the importance of C3 signaling in the activation of olfactory bulb microglia and establish this signaling pathway as necessary to maintain appropriate neuronal targeting in the olfactory system. Specifically, we showed that microglia engulf both olfactory sensory neuron axons and the axonal and dendritic terminals of tufted cells. We also reveal that complement 3 mediated signaling is necessary for activation of microglia and proper refinement of the glomerular and intrabulbar maps (Figure 6). Our work expands on previous work demonstrating that microglia shape circuits in the visual cortex (Stevens *et al.*, 2007; Tremblay *et al.*, 2010; Schafer *et al.*, 2012) and hippocampus (Paolicelli & Gross, 2011), and that much of this pruning is dependent on complement 3 signaling (Schafer *et al.*, 2012; Bahrini *et al.*, 2015; Hong *et al.*, 2016).

**Figure 6.**
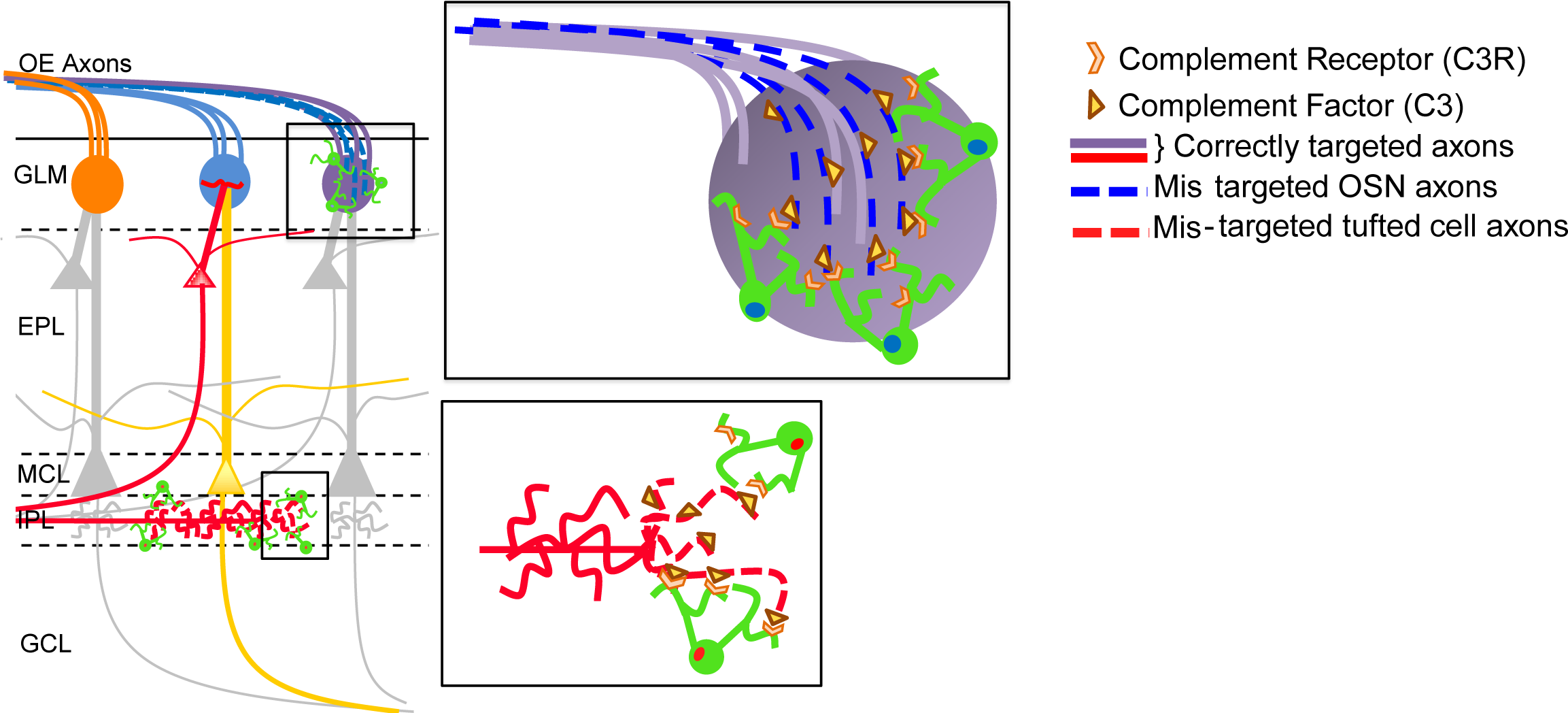
Microglia complement cascade signaling is necessary for axonal refinement in the olfactory bulb. Our model suggests that both mistargeted axons in olfactory glomeruli and axons in intrabulbar map projections are refined by microglia in a complement 3 dependent manner.

### Microglia prune glomerular circuitry in a complement 3 dependent manner

Glomerular map formation, depends both on canonical axon guidance cues (Chehrehasa *et al.*, 2005; Cho *et al.*, 2007; Rodriguez-Gil *et al.*, 2015) and developmental refinement, the mechanism of which remain unclear (Cummings & Belluscio, 2010). Our study shows that complement 3 signaling is a mechanism that regulates the refinement process as we observe a disorganized map that persist into adulthood when C3 signaling is lost. An alternate explanation for these findings is that C3 functions as an axon guidance or axon suppression cue, (Peterson *et al.*, 2017) that is independent of microglia activity. While this would explain the disorganization in the glomerular map when C3 is eliminated, it would not explain why disorganization persists when CR3, which is primarily expressed by microglia (Ehlers, 2000) is knocked out but C3 is still present. In addition, the continued presence of major glomeruli, in both conditions supports the idea that C3 loss does not cause broad axonal disorganization of OSNs. Furthermore, previous studies have shown that knocking out C3 causes a severe reduction in microglia axonal engulfment (Schafer *et al.*, 2012), supporting the idea that the disorganization we observe in the OB circuitry is due to a loss of the ability of microglia to refine mistargeted axons. Thus, our data show that C3 is necessary for glomerular map refinement and likely due to a C3 microglia dependent pruning mechanism.

This notion that axonal pruning is mediated by microglia is further supported by our observation that microglia engulf and internalize P2-GFP OSN axons in wildtype animals, showing that microglia have a normal baseline interaction with OSN axons. We observed engulfment of OSN axons around both major and minor glomeruli. However, because microglia are highly motile cells and the glomerular map is a regenerating neuronal map, so we cannot distinguish if engulfment is due to microglia pruning mistargeted axons in the minor glomeruli, or phagocytosing the continuously turning-over axons in major glomeruli. In either case, it is notable that we see engulfment in mature animals as most studies have focused on engulfment in postnatal and juvenile animals. While we cannot formally rule out the possibility that the engulfment we observe is a remnant from early development as the map is formed, previous work on the rate of microglia degradation (Peri & Nüsslein-Volhard, 2008; Schafer *et al.*, 2012; Morsch *et al.*, 2015) and the fact that our animals are four weeks beyond olfactory map maturity, suggests that the engulfed GFP particles are from recent internalization of OSN axons and that microglia are involved in the ongoing refinement of the mature glomerular map.

### C3 is necessary for the refinement of the intrabulbar map

Our examination of the intrabulbar map, specifically the lack of refinement after C3 signaling is lost, demonstrates that C3 signaling is important for the refinement of multiple, differing maps in the olfactory bulb. Unlike OSNs in the glomerular map, the tufted cells that make up the intrabulbar circuit are central projection neurons and a non-regenerating population, allowing us to ask if the observed disorganization in glomerular map following complement signaling loss requires axonal turnover—or if C3 serves a more general role in olfactory bulb circuit refinement. Our data show a failure of intrabulbar axonal refinement in both the C3(-/-) and CR3(-/-) mice indicating that C3 signaling is necessary for the developmental refinement of at least two maps in the olfactory system, and expands the role for C3 signaling in sensory map development. Interestingly the width of projections we observe in our complement knockouts are similar the width observed in both 2-week-old animals (Marks *et al.*, 2006) and animals that have been deprived of odor stimuli through naris closure (Cummings & Belluscio, 2010). This correlation could suggest that complement 3 signaling is necessary for developmental refinement or it could indicate a postnatal refinement failure due to disruption in the activation of upstream circuitry such as OSN input. We show that the glomerular map is disrupted, so an intriguing hypothesis is that the disorganization in the glomerular map has downstream effects on the intrabulbar map, and that the refinement of one map depends on refinement of the other. Notably, we observe no obvious olfactory phenotype in our C3(-/-)/CR3(-/-) animals, indicating that general olfactory function is intact. Furthermore, the germline nature of our knockout and previous studies showing the necessity of C3 signaling for map development suggest that the lack of refinement we see is due to a developmental refinement failure.

We also observe that microglia continually shape the olfactory maps in adulthood. Our data from adult animals demonstrating microglia engulfment of tufted cell synaptic terminals in both the glomerular layer and the IPL supports the idea that microglia are constantly pruning the intrabulbar projections (Cummings & Belluscio, 2008). The axonal reorganization observed in the IPL supports the theory that microglia could mediate intrabulbar map plasticity through axonal arbor pruning. Although it has been suspected that microglia actively sculpt circuits in adult animals (Nimmerjahn *et al.*, 2005; Tremblay *et al.*, 2010; Li *et al.*, 2012) most studies thus far have focused on the role of microglia in refining developmental circuits (Paolicelli *et al.*, 2011; Schafer *et al.*, 2012). Our findings, in adult animals, suggest that microglia continually refine stable olfactory circuits in adulthood and expand the repertoire of possible roles for microglial circuitry refinement in healthy, mature organisms.

### Microglia activity is dependent on C3 in the olfactory bulb

It is unclear why loss of C3 signaling results in unrefined OB maps. One possibility is that C3 signaling is necessary for microglia survival and that by disrupting the signaling we are simply eliminating microglia. However, our data show that OB microglia density persists in C3(-/-) and CR3(-/-) animals. Another explanation is that microglial function is altered when C3 signaling is disrupted. Using microglia lysosomal activity as a measure of microglial functions we observed a dramatically reduction. This demonstrates that C3 signaling is not leading to a loss of microglia, but rather it is leading to an inability of microglia to perform normal functions. While lysosomal activity is not a direct measure of engulfment ability, it has been correlated with altered synaptic pruning (Lui *et al.*, 2016). Our data further support this link and show that C3 signaling is necessary for lysosomal activity in both the glomerular and intrabulbar maps. These data and our observations of refinement failure in both maps, show C3 signaling is necessary for microglia activity which effects the OB network at multiple levels.

The plasticity of the olfactory system and the tools available for studying its formation and function make it a unique and flexible system for understanding how microglia shape neural circuits. The repeated axonal turnover associated with the glomerular map provides a rare perspective for understanding the role of microglia in circuit disruption and reorganization. Moreover, the intrabulbar map makes it possible to study microglia within the context of a flexible central circuit that does not turnover. Together, our findings lay the ground work for further studies to investigate how microglial signaling shapes olfactory map development, plasticity and stability.

## Acknowledgements

The authors would like to thank Nicholas Ryba for editing and helpful discussions.

## Author Contributions

K.S. Lehmann; Interpreted data and wrote the manuscript.

A. Wood; Designed and performed experiments and analyzed data.

D.Cummings; Designed and performed experiments and analyzed data

L. Bai; Performed experiments and analyzed data.

B. Stevens; Provided scientific support.

L. Belluscio; Conceived of study, designed experiments, interpreted data and edited manuscript

